# Structural basis for the inhibition of the SARS-CoV-2 RNA-dependent RNA polymerase by favipiravir-RTP

**DOI:** 10.1101/2020.10.21.347690

**Authors:** Katerina Naydenova, Kyle W. Muir, Long-Fei Wu, Ziguo Zhang, Francesca Coscia, Mathew J. Peet, Pablo Castro-Hartmann, Pu Qian, Kasim Sader, Kyle Dent, Dari Kimanius, John D. Sutherland, Jan Löwe, David Barford, Christopher J. Russo

## Abstract

The RNA polymerase inhibitor, favipiravir, is currently in clinical trials as a treatment for infection with SARS-CoV-2, despite limited information about the molecular basis for its activity. Here we report the structure of favipiravir ribonucleoside triphosphate (favipiravir-RTP) in complex with the SARS-CoV-2 RNA-dependent RNA polymerase (RdRp) bound to a template:primer RNA duplex, determined by electron cryomicroscopy (cryoEM) to a resolution of 2.5 Å. The structure shows clear evidence for the inhibitor at the catalytic site of the enzyme, and resolves the conformation of key side chains and ions surrounding the binding pocket. Polymerase activity assays indicate that the inhibitor is weakly incorporated into the RNA primer strand, and suppresses RNA replication in the presence of natural nucleotides. The structure reveals an unusual, non-productive binding mode of favipiravir-RTP at the catalytic site of SARS-CoV-2 RdRp which explains its low rate of incorporation into the RNA primer strand. Together, these findings inform current and future efforts to develop polymerase inhibitors for SARS coronaviruses.

## Introduction

The 2019-2020 coronavirus pandemic has spurred research into novel and existing anti-viral treatments. The SARS-CoV-2 viral RNA-dependent RNA polymerase, crucial for coronavirus replication and transcription, is a promising drug target for the treatment of COVID-19 [1]. Nucleoside analogues, including remdesivir, sofosubivir and favipiravir (T-705), have been shown to have a broad spectrum of activity against viral polymerases [2], and are being trialed against SARS-CoV-2 infections [3, 4, 5]. Recent results of the World Health Organization SOLIDARITY clinical trials indicate that remdesivir, hydroxychloroquine, lopinavir and interferon treatment regimens appeared to have little or no effect on hospitalized COVID-19 patients, as measured by overall mortality, initiation of ventilation and duration of hospital stays [6], so insight into the modes action and mechanisms of current and future drugs for COVID-19 is increasingly important. The structures of the SARS-CoV RdRp [7], the SARS-CoV-2 RdRp [8], the SARS-CoV-2 polymerase bound to an RNA duplex [9], to RNA and remdesivir [10, 11], and to the helicase nsp13 [12] were all recently determined by electron cryomicroscopy (cryoEM). The suggested modes of action of favipiravir, a purine nucleic acid analogue derived from pyrazine carboxamide (6-fluoro-3-hydroxy-2-pyrazinecarboxamide), against coronaviruses comprises nonobligate chain termination, slowed RNA synthesis, and mutagenesis of the viral genome [13]. Here we report the structure of the SARS-CoV-2 RdRp, comprising subunits nsp7, nsp8 and nsp12, in complex with template:primer double-stranded RNA and favipiravir ribonucleoside triphosphate (favipiravir-RTP), determined by cryoEM at 2.5 Å resolution.

## Materials and methods

### General reagents

Favipiravir (T-705-RTP) was purchased from Santa Cruz; upon receipt a stock solution of 50 mM T-705-RTP resuspended in 100 mM of Tris, pH 7.5 buffer was aliquoted and stored at −80° C. Phosphoramidites were purchased from Sigma-Aldrich or Link Technologies and used without further purification. Acetonitrile and other reagents (cap solutions, de-block solution and oxidizer solution) were purchased from Sigma-Aldrich. Primer Support 5G for A, G, C or U (with loading at ~300 μmol/g) respectively, was purchased from GE Healthcare.

### Spectroscopy of chemical compounds

Mass spectra were acquired on an Agilent 1200 LC-MS system equipped with an electrospray ionization (ESI) source and a 6130 quadrupole spectrometer (LC solvents: A, 0.2% formic acid in H_2_O and B, 0.2% formic acid in acetonitrile). ^31^P-Nuclear magnetic resonance (^31^P-NMR) spectra of the T-705-RTP stock solution were acquired using a Bruker Ultrashield 400 Plus operating at 162 MHz.

### Protein purification

The gene-optimized cassette (GeneArts/Thermo Fisher) encoding a SARS-CoV-2 RdRp polyprotein, codon optimized for *Spodoptera frugiperda*, containing nsp5-nsp7-nsp8-nsp12 was cloned, with a C-terminal TEV-double StrepII tag, into a pU1 vector for a modified MultiBac baculovirus/insect cell expression [14]. The cell pellet was lysed in a buffer of 50 mM Tris-HCl (pH 8.0), 250 mM NaCl, 2 mM MgCl_2_, 1 mM DTT, and loaded onto a Strep-Tactin column (Qiagen). The protein complex was eluted in lysis buffer supplemented with 5 mM desthiobiotin. The RdRp was further purified by size exclusion chromatography on a Superdex 200 16/60 size-exclusion column (Cytiva) in a buffer of 20 mM HEPES (pH 7.5), 200 mM NaCl, 2 mM MgCl_2_ and 1 mM TCEP. Fractions containing nsp7-nsp8-nsp12 were concentrated, flash-frozen in liquid nitrogen, and stored at −80°C as single-use aliquots.

Nsp7 and nsp8 were cloned into the NcoI-NotI, and BamHI-NotI sites of pAcycDuet1 (Millipore), and a pET28 vector modified to encode an N-terminal 6x-His-Sumo tag in frame with the multiple policloning site. Protein expression was pursued in *Escherichia coli* (*E. coli*) BL21(DE3) Star cells co-transformed with the nsp7 and nsp8 plasmids. Cells were grown in ZY auto-induction media [15] at 37°C, until an optical density at 600 nm of 0.8 was attained, at which point the temperature was reduced to 18°C, and the incubation was continued overnight. Cells were lysed by sonication in a buffer containing 500 mM NaCl, 20 mM HEPES pH 7.5, 5 mM benzamidine, 20 mM imidazole, 1 mM TCEP. Proteins were captured with Co^2+^-conjugated-IMAC resin (Cytvia), eluted in a buffer containing 300 mM NaCl, 20 mM HEPES pH 7.5, 300 mM imidazole, 1 mM TCEP, followed by overnight cleavage of the 6x-His-Sumo tag by Ulp1, and subjected to subtractive IMAC to remove cleaved tags, uncleaved nsp8, and Ulp1. The nsp7-nsp8-containing solution was then diluted to 150 mM NaCl, further purified via cation-exchange, and size exclusion chromatography in assembly buffer (100 mM NaCl, 20 mM HEPES pH 7.5, 2 mM MgCl_2_, and 1 mM TCEP; Superdex 200 16/60, Cytiva). Fractions containing nsp7-nsp8 were concentrated, flash-frozen in liquid nitrogen, and stored at −80°C as singleuse aliquots.

### RNA synthesis

The RNA template, 5’-rUrUrUrUrUrCrArUrArArCrUrUrArArUrCrUrCrAr CrArUrArGrCrArCrUrG-3’, and RNA primer 5’-rCrArGrUrGrCrUrArUrGrUr GrArGrArUrUrArArGrUrUrArU-3’ were prepared by solid phase synthesis on an ÄKTA oligopilot plus 10 (GE Healthcare). RNAs were cleaved from the solid support by treating with 4 mL of a 1:1 mixture of 28% wt NH_3_/H_2_O solution and 33% wt CH_3_NH_2_/EtOH solution at 55°C for 30 minutes, then the silyl protecting groups were removed by treating with 3 mL of 1:1 mixture of triethylamine trihydrofluoride and DMSO at 55°C for 90 minutes. Then, 30 mL of cold 50 mM NaClO_4_ in acetone was added to precipitate the RNA product. After centrifugation, the pellet of RNA was dissolved in 5 mL of water and passed through a Sep-Pack C18 Cartridge, 5 g sorbent (Waters). Eluates containing RNA were combined and lyophilized. Mass spectroscopy of the template RNA, found m/z ([M-7H^+^]) = 1341.8 (theoretical 1342.1), found m/z ([M-6H^+^]) = 1565.6 (theoretical 1565.9); and of the primer RNA, found m/z ([M-6H^+^]) = 1278.6 (theoretical 1278.9), found m/z ([M-5H^+^]) = 1534.5 (theoretical 1534.9).

### Annealing of primer:template RNA duplexes

Single-stranded primer and template RNAs (Fig. 1A) were resuspended in deionized and purified H_2_O to a final concentration of 200 μM, mixed at an equimolar ratio, and incubated for 5 minutes in a heat block at 95°C in Eppendorf tubes with punctured lids. The heat block was then removed from the heating device and allowed to cool to ambient temperature. Annealed dsRNAs were then dispensed as single-use aliquots, flash frozen in liquid nitrogen, and stored at −80°C.

**Figure 1:**
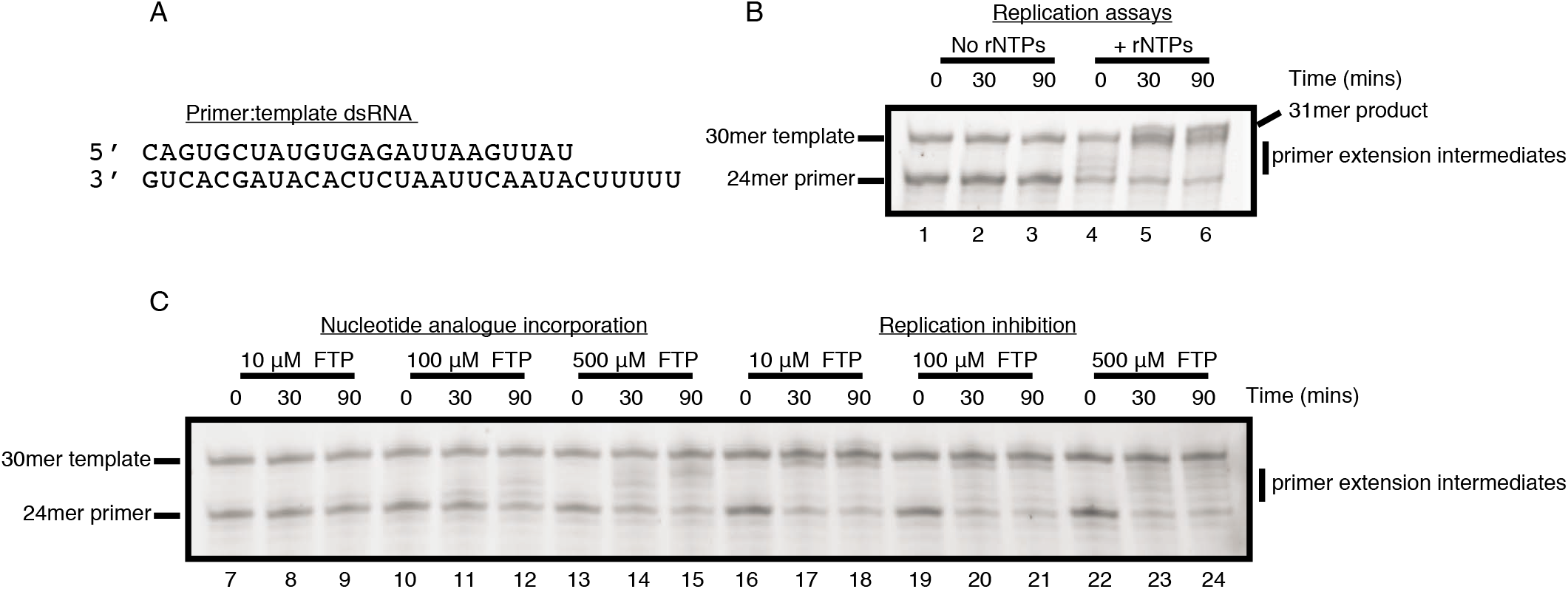
Reconstitution of SARS-CoV-2 RdRp activity and inhibition by favipiravir-RTP (FTP). (**A**) Sequence of the annealed primer (top):template (bottom) dsRNA duplex employed in biochemical assays. (**B**) Reconstituted SARS-CoV-2 RdRp extends the 24mer primer in the presence (lanes 4-6), but not the absence (lanes 1-3), of ribonucleotides (rNTPs). A 31mer product is present in lane 6 due to addition of a non-templated base to the primer strand. (**C**) Favipiravir is weakly incorporated into the 24mer primer strand (lanes 7-15), and suppresses RNA replication by the SARS-CoV-2 RdRp.

### Assembly of the apo-RdRp complex

Purified apo-RdRp, assembled as above, was mixed immediately after purification with annealed dsRNA, resuspended in ddH_2_O, to a final protein:RNA concentration of 8:20 μM, and was incubated at room temperature for 5 minutes prior to addition of favipiravir-RTP to a final concentration of 100 μM. Thereafter, complexes were incubated for an additional 30 minutes at room temperature, prior to another round of centrifugation at 4°C, transferred to fresh Eppendorf tubes, and used immediately for the preparation of cryoEM grids.

### Primer extension, drug incorporation, and replication inhibition assays

All assays (Fig. 1 B-C) were performed at room temperature in assembly buffer. Reaction conditions were designed to approximate as closely as possible those employed in production of vitrified EM grids. For primer extension assays, 6 μM RdRp (prepared as above) was assembled with 6 μM dsRNA, and mixed with rATP/rGTP (at a final concentration of 500 μM). For drug incorporation assays, RdRp:dsRNA complexes were assembled, mixed with favipiravir-RTP (at final concentrations of 10, 100, and 500 μM), and reactions allowed to proceed. For replication inhibition assays, RdRp:dsRNA complexes were assembled, preincubated with favipiravir-RTP (at final concentrations of 10, 100, and 500 μM) for a period of 30 minutes, prior to addition of rATP/rGTP to a final concentration of 500 μM. Samples were removed at the indicated time points (0, 30, and 90 minutes); the reactions were stopped with a 1:1 addition of quenching buffer (98% formamide, 10 mM EDTA), flash frozen and stored at −21°C prior to gel analyses.

### Specimen analysis

20% polyacrylamide, 8 M urea gels (0.75 mm thick, 20 cm long) were run at 15 W in TBE buffer for 2 hours. The RNA gel was stained using SYBR Gold Nucleic Acid Gel Stain (Invitrogen). Fluorescence imaging was performed using an Amersham Typhoon imager (GE Healthcare) and quantified using Image Quant TL software (version 7.0).

### CryoEM and atomic model building

The complex, prepared as described above, was used at a concentration of 1.3 mg/mL for preparing cryoEM grids. All-gold HexAuFoil grids with a hexagonal array of 280 nm diameter holes, 700 nm hole-to-hole spacing, and 330 Å foil thickness were made in-house, and plasma treated as described in [16]. Graphene was grown by chemical vapor deposition, transferred onto allgold UltrAuFoil R0.6/1 grids (QuantiFoil), and partially hydrogenated following a previously described procedure [17]. Grids were rapidly cooled using a manual plunger [18] in a 4°C cold room. A3 μL volume of the protein solution was pipetted onto the foil side of the grid and then blotted from the same side with filter paper (Whatman No. 1) for 11-14 seconds. The grids were immediately plunged into liquid ethane, kept at 93K in a cryostat [19], and were stored in liquid nitrogen until they were imaged in the electron cryomicroscope.

We acquired electron micrographs from four grids: three HexAuFoil grids (total 52,881 multiframe micrographs) and one partially hydrogenated graphene-coated grid (11,096 multiframe micrographs) (Fig. 2A, Table S1). Other surfaces that were screened during initial specimen preparation included graphene functionalized with amylamine or hexanoic acid and graphene oxide, none of which improved the orientation distribution or yielded 2D classes indicating degradation of the complex more severe than due to the air-water interface alone. Thus, they were not included in the dataset for high-resolution reconstruction. The high density of foil holes on the HexAuFoil grid, combined with fast imaging by aberration-free image shift and a fast, direct-electron detector (Falcon 4, 240 Hz) allowed us to acquire more than 200 micrographs per hour for 14 consecutive days. The number of micrographs that can be acquired from a single HexAuFoil grid is limited only by the ice contamination rate in the vacuum of the electron microscope, rather than by the number of holes on one grid, even with the fastest available data collection setups.

**Figure 2:**
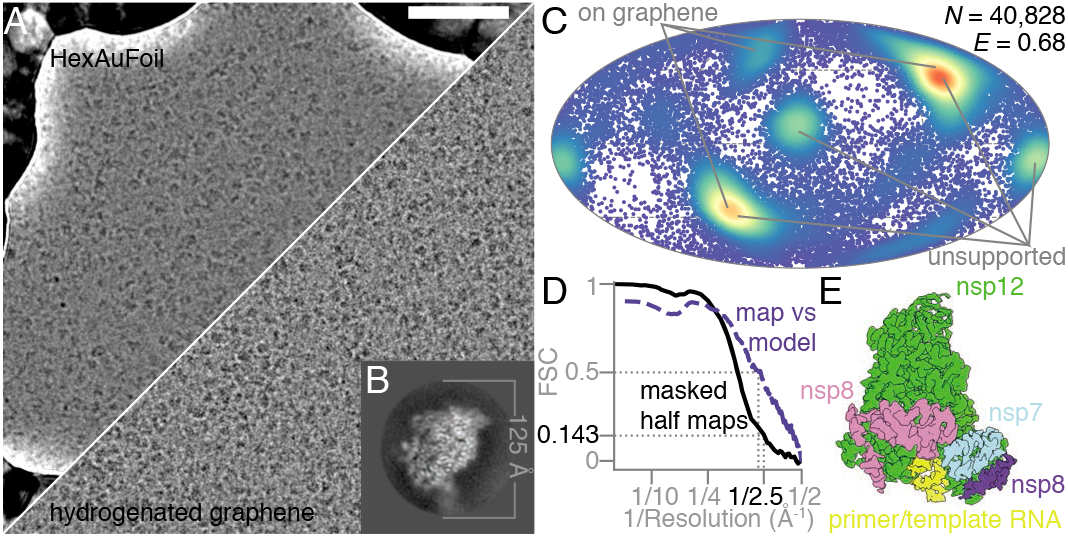
Electron cryomicroscopy of RdRp complexes in the presence of RNA and favipiravir-RTP. (**A**) Electron cryomicrographs of the reconstituted complexes in unsupported ice (HexAuFoil grid, upper), and on hydrogenated graphene (lower) were used for structure determination. The scale bar is 500 Å. (**B**) This 2D class average from the images of the complex, containing RNA, corresponds to the most frequent orientation of the particles in the thin film of vitreous water. (**C**) The orientation distribution, with efficiency, *E_od_* [23], of the particles used in the reconstruction is plotted on a Mollweide projection, with the most common views on each type of grids marked with gray lines. (**D**) The Fourier shell correlation (FSC) between the two independent masked half maps, and between the final map and the atomic model, is plotted versus resolution. (**E**) An overview of the EM map of the polymerase complex is colored by subunit: green - nsp12, blue - nsp7, pink and purple - two copies of nsp8, yellow - template:primer RNA.

All data were processed in RELION 3.1 [20], using particle picks imported from crYOLO 1.5 [21] (Fig. S1) with manual retraining of the model. After four rounds of 2D classification, it was clear that the specimen adopted a preferred orientation (Fig. 2B), as previously reported [9]. The uniformity of the particle orientation distribution, and the isotropy of the final 3D reconstruction, were improved by combining data from HexAuFoil and partially hydrogenated graphene-coated UltrAuFoil EM grids (Fig. 2C). After three rounds of 3D classification, we obtained a 2.5 Å resolution isotropic map of the full complex from 40,828 particles (Fig. 2D-E). A different set of 140,639 particles produced a 2.5 Å resolution map of the nsp12 subunit alone. The remaining 99% of the particles were discarded, as these corresponded to either the over-represented view of the nsp12 subunit alone, or to structurally heterogeneous complexes that did not align to high resolution.

Optical aberrations were refined per grid, the astigmatism per micrograph, and the defocus per particle. Particle movement during irradiation was tracked separately for the graphene and the HexAuFoil datasets, using Bayesian polishing [22]. The dataset on graphene shows typical beam induced motion at the onset of irradiation, whereas this movement is eliminated by the use of the HexAuFoil grids, which provided superior data quality, especially at the onset of irradiation, when the specimen is least damaged. Of the particles contributing to the final 3D refinement, 80% originated from the HexAuFoil grids, and 20% from the partially hydrogenated graphene grid. This was necessary to improve the particle orientation distribution: the efficiency, *E_od_*, increased from 0.54 in unsupported ice, and 0.57 on partially hydrogenated graphene to 0.68 using a combination of particles from both support surfaces (Fig. 2C) [23]. External reconstruction in SIDESPLITTER was performed [24]. The atomic model was built based on a previously published atomic model of the nsp12-nsp7-nsp8 complex bound to RNA and remdesivir-RTP (PDB-7BV2) [10], with manual model building in Coot [25, 26], and real-space refinement in Phenix [27].

## Results

To confirm the integrity of the assembled nsp7-nsp8-nsp12 SARS-CoV-2 RdRp complexes, we first performed primer extension activity assays. Consistent with prior publications, we observed efficient primer extension in the presence of natural ribonucleotide triphosphates (rNTPs) (Fig. 1A-B: lanes 4-6) [8, 9, 10, 12, 11]. Having verified the activity of the reconstituted RdRp, we then investigated the ability of the enzyme to incorporate favipiravir-RTP into the primer strand. In contrast to natural rNTPs, favipiravir-RTP appears to be a comparably poor substrate, with a low fraction of primer extension observed over the range of experimental conditions tested (Fig. 1C: lanes 7-15). The formation of larger RNA products observed at later time-points (in particular lanes 12 and 15), must arise through promiscuous base-pairing with both uracil and cytosine in the template strand, and is consistent with the reported mutagenicity of favipiravir towards viral genomic RNA in vivo [28, 29, 13]. Finally, we tested whether favipiravir-RTP can inhibit primer extension in the presence of rNTPs. Despite the low incorporation of favipiravir-RTP observed in primer extension assays, it nevertheless suppresses completion of RNA replication at all concentrations tested, even when rNTPs are present at a considerable excess over the inhibitor (Fig. 1C: lanes 16-24). These assays are thus consistent with a previously proposed mode of action of favipiravir-RTP, comprising both slowed RNA synthesis and non-obligate chain termination [13].

To investigate the structural basis of this activity, we determined the structure of the nsp7-nsp8-nsp12 SARS-CoV-2 RdRp complex, in the presence of template:primer RNA and favipiravir-RTP, by cryoEM (Fig. 2, Fig. S1, Fig. S2, Table S1). The overall structure of the polymerase complex observed (Fig. 3A, Fig. S3), comprising one nsp12, one nsp7, two nsp8 subunits, and a template:primer RNA, is nearly identical to those previously described [10, 9]. With 2.5 Å resolution, and the reduced effects of radiation damage afforded by movement-free imaging [16], the map we determined is suitable for unambiguous atomic model building. We resolved and modelled some additional density in the N-terminal nidovirus RdRp-associated nucleotidyltransferase (NiRAN) domain of nsp12, including a pyrophosphate and a Mg^2+^ ion (Fig. 3B). These are surrounded by residues which are conserved between SARS-CoV and SARS-CoV-2, including K73, R116, D218 of the active site of the NiRAN domain [12]. However, we did not find density for a possible nucleotide linked to the pyrophosphate at this site. A previous study [11] also observed density consistent with a pyrophosphate, only in a pre-translocated RdRP-RNA complex, but did not attribute it due to insufficient resolution.

**Figure 3:**
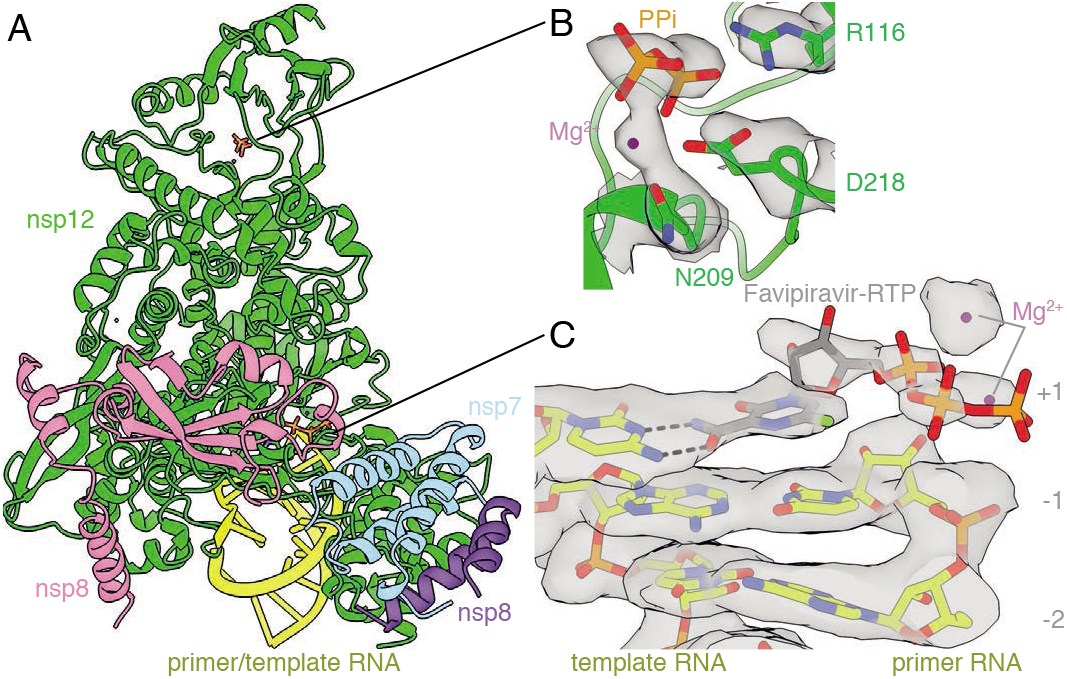
Structure of the RdRp complex bound to dsRNA and favipiravir-RTP. (**A**) The overall structure of the complex is shown, colored by subunit, in the same way as in Fig. 2E. The black lines point to the approximate positions of the catalytic sites, displayed in panels (**B**) and (**C**), where the coloring is by heteroatom, and the EM map is contoured in gray. (**B**) A pyrophosphate is present at the NiRAN catalytic site of the enzyme, and is coordinated by key conserved residues in this domain. (**C**) The template:primer RNA duplex is shown, along with favipiravir at the catalytic site (+1) of the polymerase. Hydrogen bonds are indicated.

We found that favipiravir and two catalytic Mg^2+^ ions are present at the catalytic site of the complex (Fig. 3C, Fig. S2). Favipiravir is stacked onto the 3’ nucleotide of the primer strand, and forms a non-canonical base pair, through its amide group, with the displayed cytidine in the template RNA strand, consistent with favipiravir mimicking a guanosine base. The weaker density for the inhibitor, compared to that for the neighboring RNA bases, indicates partial occupancy of the site. The density is most consistent with the inhibitor binding non-covalently to the polymerase, with little to no covalent incorporation. The biochemical assays indicated only minimal incorporation of favipiravir-RTP into the primer under the conditions used to prepare the complex for structural analysis (Fig. 1; lane 11). Density for the *α* and *β* phosphates, which are coordinated by two neighboring magnesium ions, can be clearly traced. The density for the favipiravir base appears discontinuous from the density for the ribose bound to it; however, we verified using spectroscopy that the glycosidic bond was stable in conditions used to prepare the complex for cryoEM (Fig. S4). We also observed a weak signal, adjacent to the trisphosphate moiety of favipiravir-RTP, consistent with a pyrophosphate molecule. ^31^P-NMR analysis revealed that the supplied favipiravir-RTP contained pyrophosphate contaminant at approximately equivalent amounts to the favipiravir-RTP inhibitor (Fig. S5). We show that pyrophosphate does not inhibit RdRp activity in the presence of rNTPs (Fig. S6). Pyrophosphate was modelled at this site in the SARS-CoV-2 RdRp nsp12-RNA-remdesivir complex (PDB ID 7BV2) [10]. However, in the influenza RNA polymerase this site is occupied by the trisphosphate of the incoming rNTP (PDB 6SZV [30]). In contrast, the current structure indicates that favipiravir-RTP is bound at the RdRp catalytic site in a non-productive configuration, as discussed below.

We observe signal for the RNA base pairs as synthesized, indicating no translocation of the RNA has occurred in the majority of the intact complexes during the incubation period of 30 minutes. This supports the interpretation that favipiravir stalls replication by non-covalent interactions at the active site, rather than by covalent incorporation into the replicating strand. Neither base stacking nor base pairing distances in the presence of favipiravir differ from those expected for natural nucleotides. In addition to base-pairing with cytosine from the template strand, favipiravir is coordinated by Lysine 545 in the F1 domain of nsp12, which is positioned to accept hydrogen bonds from the nitrogen atom in the pyrazine ring or donate to the fluorine atom of the inhibitor, although in both instances the distances are long (3.4 A and 3.7 Å, respectively), suggesting possible water-mediated contacts. These interactions are potentially functionally important because an arginine substitution of the equivalent residues in the chikungunya virus (CHIKV) and influenza virus H1N1 RdRp’s is responsible for their decreased susceptibility to favipiravir [2, 31]. Two nearby arginines (R553, R555) in the nucleoside entry channel are flexible, with no signal visible for the guanidinium group of R555. The 2’ hydroxyl of the favipiravir nucleotide analogue forms a hydrogen bond to residue N691 of nsp12. A serine (S682) is positioned, as previously hypothesized [9], to play a role in coordinating the nucleotide for incorporation. The hydrophobic valine (V557) is stacked against the +1 base in the RNA template.

## Discussion

In this study, we determined the cryoEM structure of favipiravir-RTP at the catalytic site of the SARS-CoV-2 RdRp, in complex with template:primer dsRNA, and investigated the influence of this nucleotide analogue inhibitor on RNA synthesis in vitro. We observed that favipiravir-RTP is an inefficient substrate for the viral RdRp in primer extension assays, and propose that this is a consequence of a catalytically unproductive conformation adopted by the drug in the polymerase active site. The non-productive binding mode of favipiravir-RTP to the catalytic site of RdRp observed here explains the inefficient rate of covalent incorporation in primer extension assays. In the binding mode reported here, the *β*-phosphate of favipiravir-RTP is not aligned for in-line nucleophilic attack by the 3’OH of the P–1 nucleotide. An optimal geometry would require rotation of the ribose O5’-*α*P bond by 120°. The structure of a productive influenza polymerase RNA-rNTP complex (PDB-6SZV) [30] showed that the triphosphates of the P+1 nucleotide engage the site possibly occupied by pyrophosphate (the product of rNTP incorporation) in the structure reported here (Fig. 4, Fig. S3). One might hypothesize that, pyrophosphate contributes to the slow rate of favipiravir-RTP incorporation by directing the triphosphate moiety of favipiravir-RTP into a non-productive binding mode. However, pyrophosphate inhibition assays (Fig. S6) indicate that pyrophosphate at concentrations between 10 and 500 μM does not affect the replication activity of RdRp. We therefore suggest that the non-productive configuration of favipiravir-RTP is promoted by an intrinsic feature of the favipiravir moiety. One possibility is that water-mediated hydrogen bonds linking the fluorine atom of the pyrazine ring to the oxygen atoms of the *β*-phosphate group to reconfigure the triphosphate into a non-productive position, although we do not unambiguously identify water molecules bound to favipiravir-RTP in the map at this resolution. In support of this mechanism, T-1105-RTP, which lacks the fluorine, is a more efficient substrate than favipiravir-RTP for SARS-CoV-2 RdRp [13].

**Figure 4:**
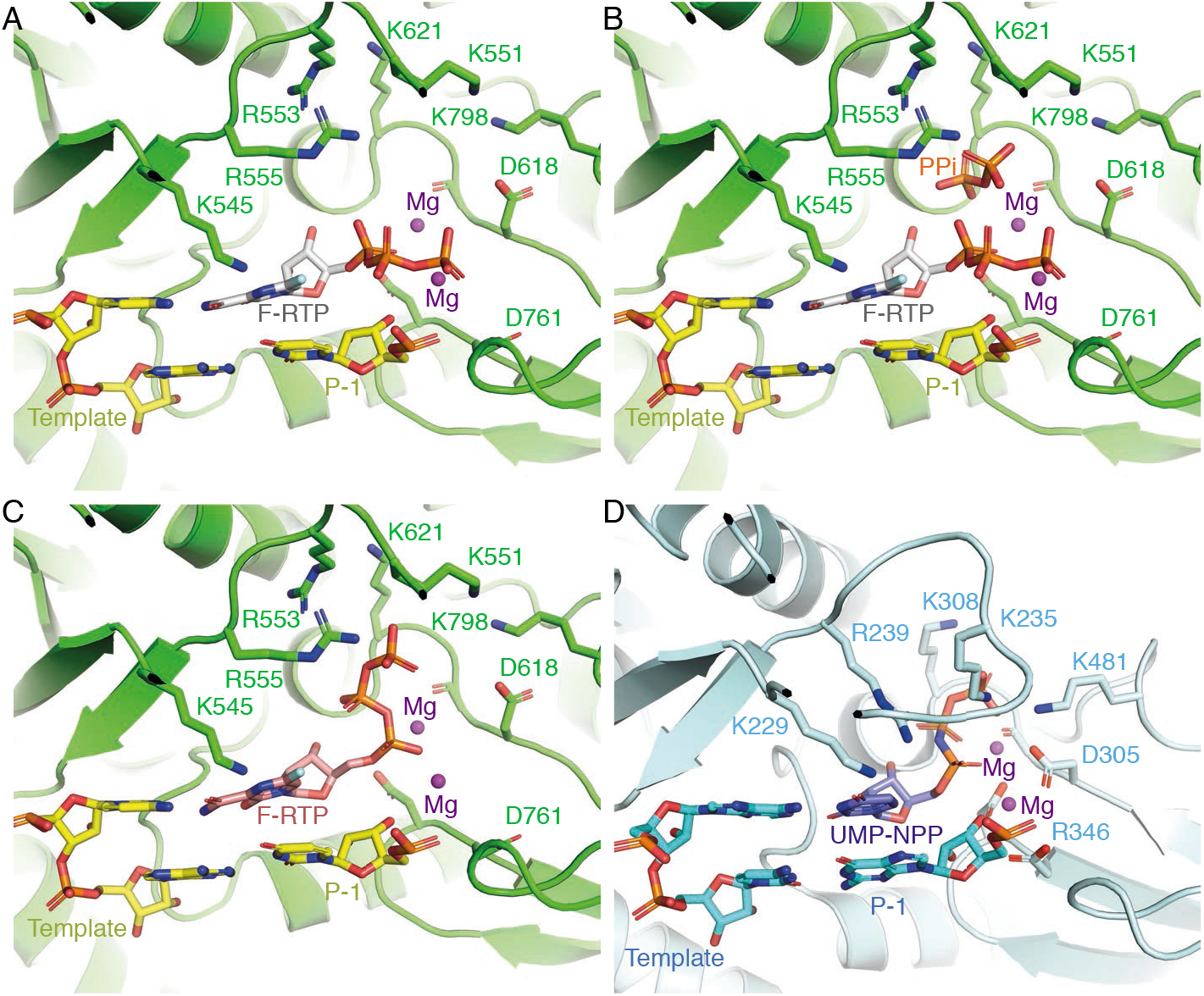
Coordination of favipiravir-RTP in the active site of the SARS-CoV-2 RdRp. (**A**) A non-productive conformation of favipiravir-RTP (F-RTP), base-paired to the P+1 nucleotide of the template strand, as shown, was observed in the cryoEM density map. For clarity, the primer strand is omitted and only the P+1 and P – 1 nucleotides of the template strand are shown. (**B**) The non-productive conformation may be favored in the presence of pyrophosphate (PPi), which is a possible by-product from the incorporation of rNTPs into the RNA. (**C**) A hypothetical productive conformation of favipiravir-RTP, which may lead to its incorporation into the primer RNA strand, is modelled. (**D**) For comparison, the position of an incoming non-hydrolyzable UTP analog in the bat influenza polymerase elongation complex is shown (PDB-6SZV) [30].

While this manuscript was in preparation, another cryoEM reconstruction and an associated atomic structure of the RdRP:dsRNA:favipiravir-RTP complex became publicly available (EMD-30469, PDB 7CTT) [32]. This map is distorted along one direction, an artefact characteristic of anisotropic filling of Fourier space in the reconstruction. Multiple factors, including complex dissociation, denaturation at the air water interface, and preferred orientation, also impeded our initial attempts for high-resolution structure determination of this complex. We managed to circumvent these by acquiring a large cryoEM dataset and using a combination of HexAuFoil and partially hydrogenated graphene-coated UltrAuFoil EM grids. We make the complete dataset publicly available via the Electron Microscopy Public Image Archive (accession number 10517) in the hope that it will be useful for software development for handling large data sets of structurally heterogeneous cryoEM specimens.

For now, cryoEM structure determination of this RdRp-RNA complex in combination with different inhibitors remains far from a routine task. Still, this and other published structures of the complex will aid rational drug design, and, in the interim, biochemical optimization of the stability of the complex might help yield a specimen that is more stable in the thin layers of buffer used for cryoEM. This would then make it suitable for rapid structure determination amongst an array of candidate compounds. The need for a derivatized graphene surface to overcome the strong interactions with the air-water interfaces highlights the problem that surfaces present in cryoEM specimen preparation; further improvements in the specimen preparation process are still needed to enable high throughput for drug design and analysis.

The structure reported here suggests at least three possible directions for further efforts towards drug design against the SARS-CoV-2 RdRp. The first is modification of the favipiravir-RTP to bring the *α*-phosphate into a more favorable confirmation for covalent incorporation at the terminus of the extending strand, leading to delayed chain-termination and/or more efficient mutagenesis of the viral genome [13]. Second, analogues of pyrophosphate bound to the RdRp active site, combined with catalytically inefficient rNTP mimics, might promote a stalled state of the polymerase. Third, modifying the fluorine position or adding other functionalities could make it better hydrogen bond acceptor. High resolution structure determination may aid in these directions as new compounds are developed and tested as treatments for COVID-19.

## Data availability

The electron scattering potential map is deposited in the Electron Microscopy Data Bank with accession code EMD-11692 and the atomic model is deposited in the Protein Data Bank under accession code PDB ID 7AAP. The complete, unprocessed cryoEM dataset is deposited in the Electron Microscopy Public Image Archive under code EMPIAR-10517.

## Acknowledgements

We thank P. Midgley for enabling access to the Department of Materials Science & Metallurgy of the University of Cambridge for data collection during the pandemic, and S. Scheres for helpful advice during data processing. We also thank J. Grimmett and T. Darling of the LMB Scientific Computing and are grateful to S. Chen, G. Cannone, G. Sharov, A. Yeates and B. Ahsan of the LMB Electron Microscopy Facility for technical assistance and enabling access to the microscopy facility during the pandemic. This work was supported by a Vice-Chancellor’s Award (Cambridge Commonwealth, European and International Trust) and a Bradfield scholarship (to KN), and Medical Research Council grants MC_UP_120117 (CJR), MC_UP_1201/6 (DB), MRC U105184326 and Wellcome 202754/Z/16/Z (JL), MC_UP_A024_1009 (JS), and Cancer Research UK grant C576/A14109.

## Supplementary information

**Figure S1:**
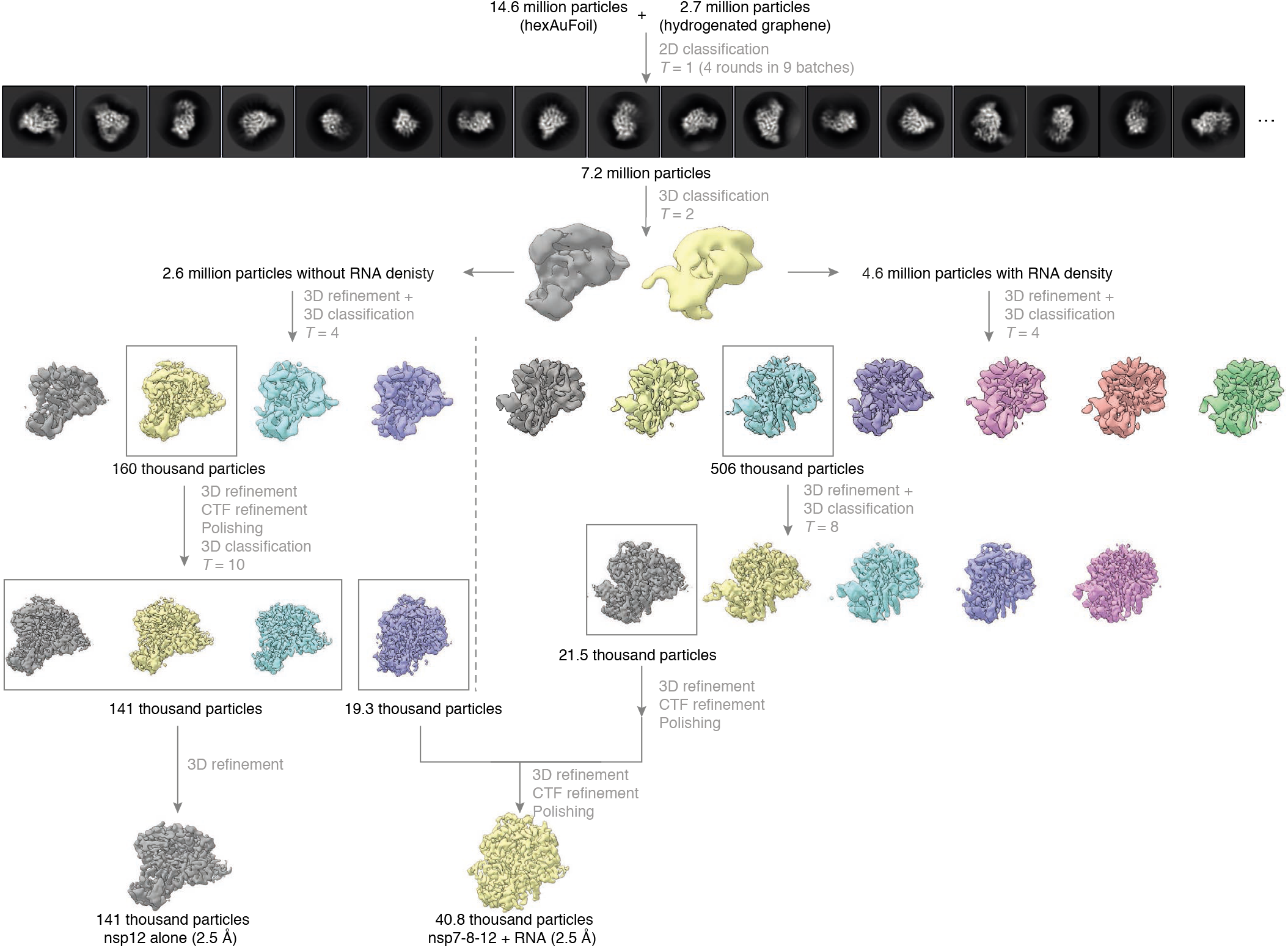
Cryo-EM data processing summary. The flowchart shows the main steps in the data processing, from particle picking, through classification, to final maps. A selected subset of the initial reference-free 2D class averages and all the intermediate 3D class averages computed during the processing of this dataset are shown. All 3D class averages, selected for subsequent rounds of processing, are boxed in gray, and the number of particles in each of these is shown. Further attempts to process the discarded classes are omitted from this chart for clarity, as these data did not contribute to the final particle set.

**Figure S2:**
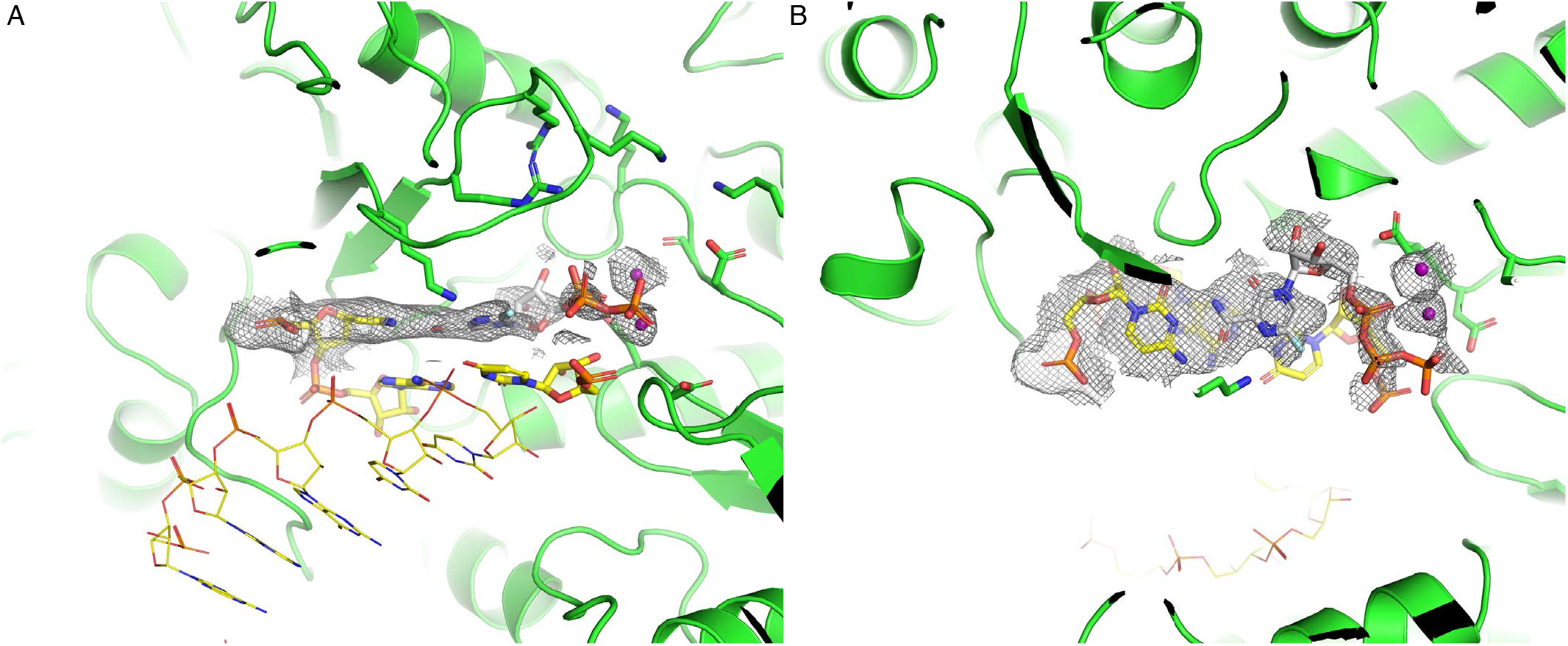
CryoEM density around the catalytic site of the SARS-CoV-2 RdRp suggests a non-productive binding mode of Favipiravir-RTP. Two orthogonal views of the polymerase active site, with map density contoured around favipiravir-RTP and Mg^2+^ (**A-B**) are shown.

**Figure S3:**
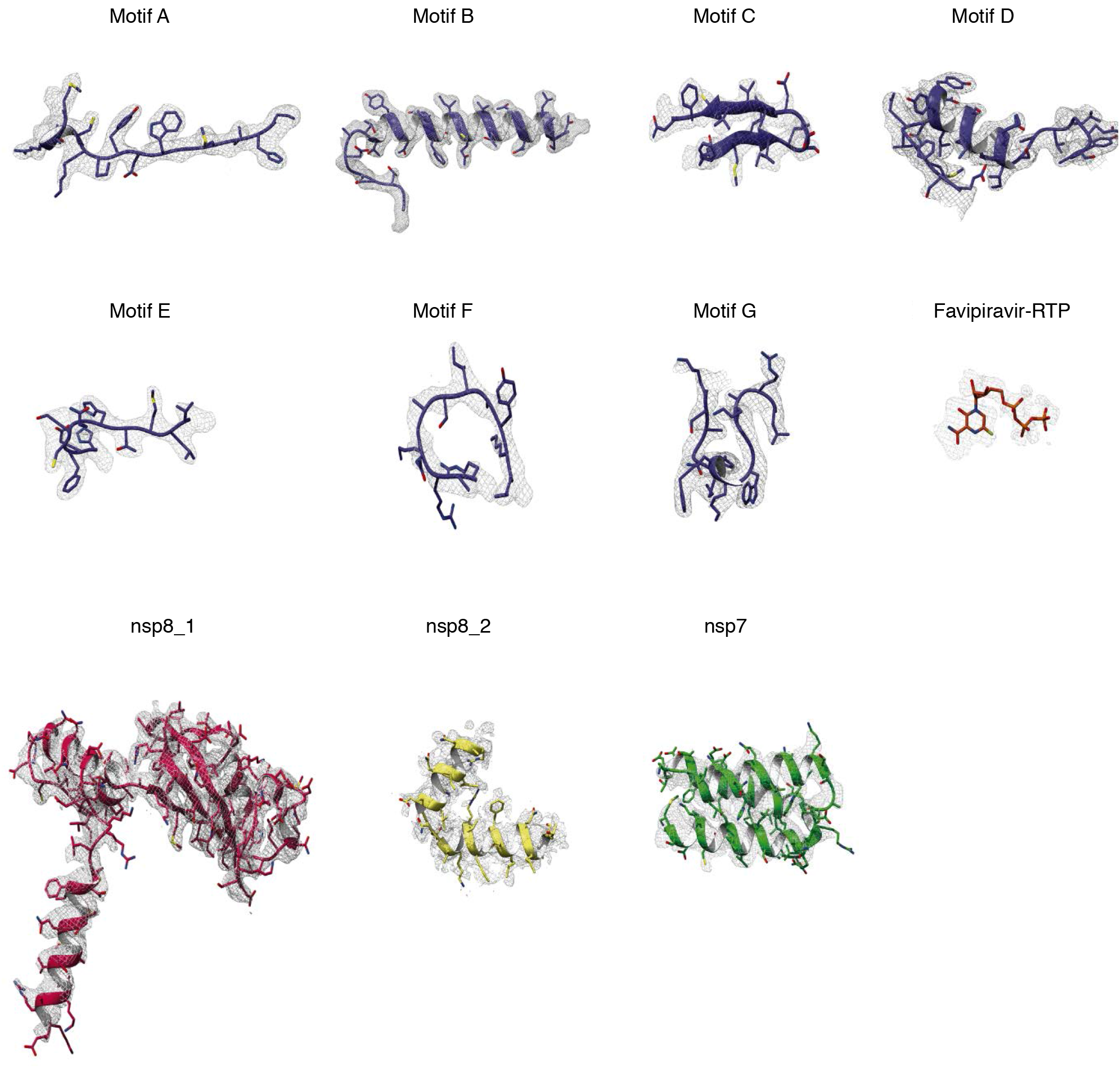
Map quality and model fitting. Map density corresponding to the indicated catalytic motifs of the nsp12 subunit, favipiravir-RTP, and nsp8_1, nsp8_2, and nsp7 subunits, contoured around the structure of the SARS-CoV-2 RdRp:dsRNA:favipiravir-RTP complex.

**Figure S4:**
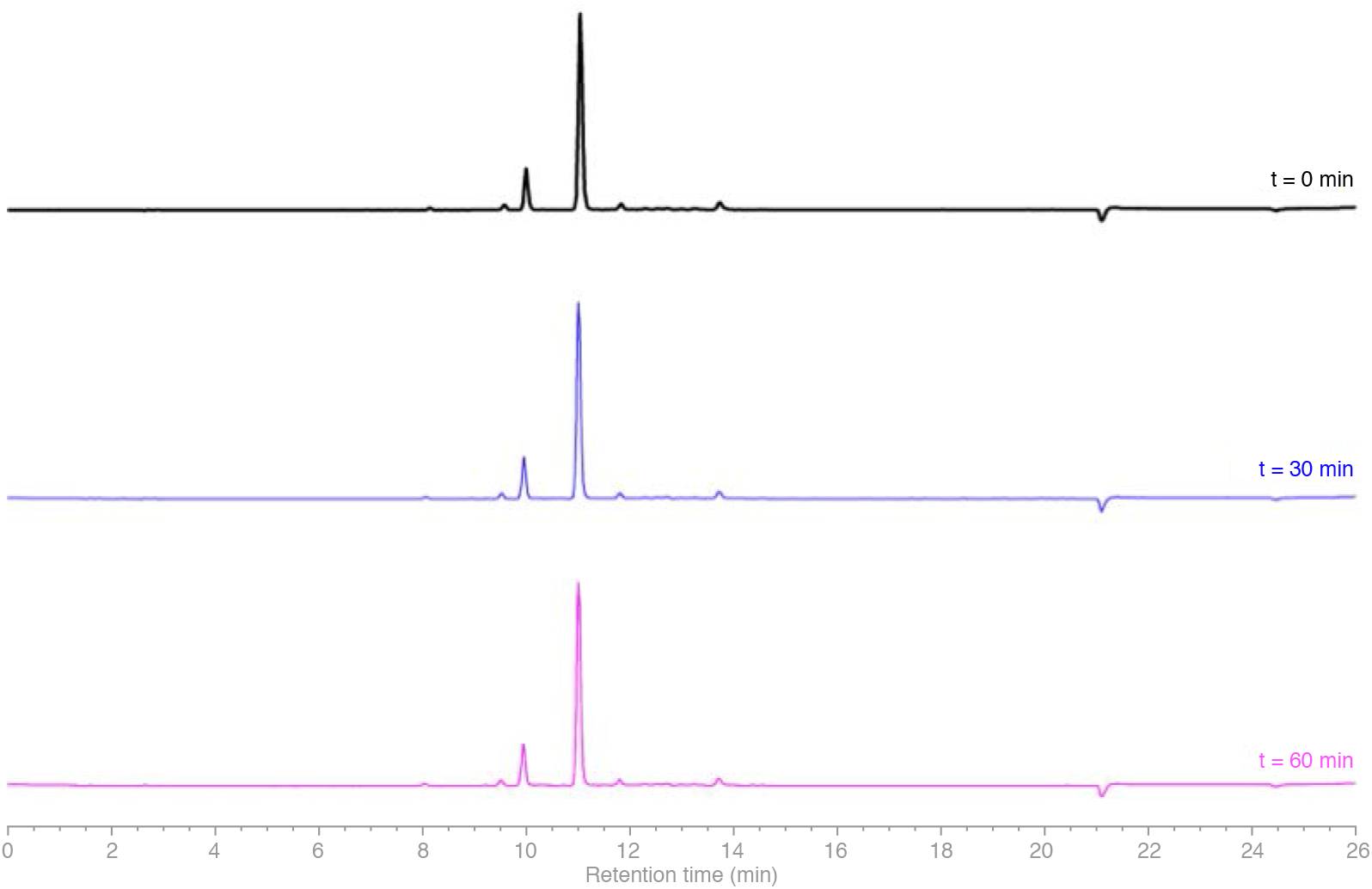
HPLC stability assay of T-705-RTP. The stability of T-705-RTP (0.5 mM) under enzymatic assay conditions was monitored using HPLC, via UV absorbance at 360 nm. The solution was incubated at 20°C and was injected into the HPLC device the indicated certain time points. No obvious changes in the peaks were observed after 1 hour.

**Figure S5:**
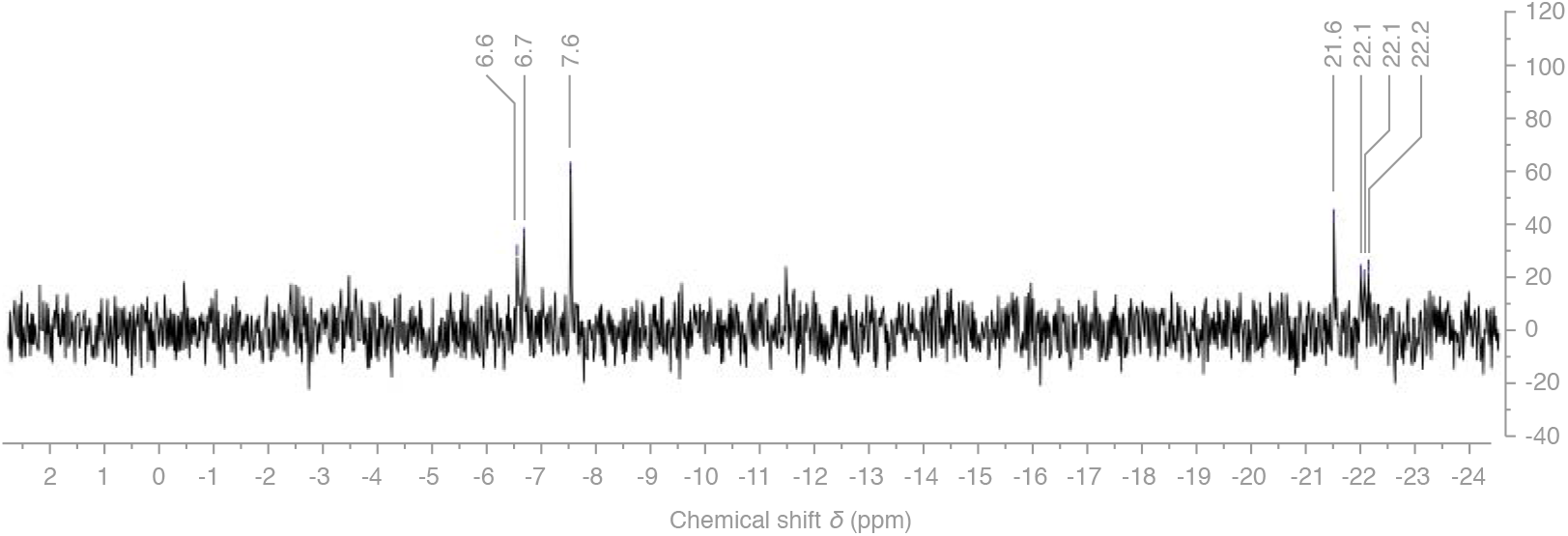
^31^P-NMR spectrum of T-705-RTP (0.5 mM) in D2O/H2O = 9:1 containing 1 mM Tris buffer at pH 7.5. The doublet at *δ* = −6.7 ppm was assigned to one of the *α*- or *γ*-phosphates of the triphosphate moiety, the other expected doublet might have been broadened by chelation to traces of paramagnetic metal ions. The triplet at *δ* = −22.1 ppm was assigned to the *β*-phosphate of the triphosphate moiety. The singlet with *δ* = −7.6 ppm is assigned to inorganic pyrophosphate, a potential contaminant associated with the process of synthesizing T-705-RTP. The singlet with *δ* = −21.6 ppm is tentatively assigned to cyclic trimetaphosphate.

**Figure S6:**
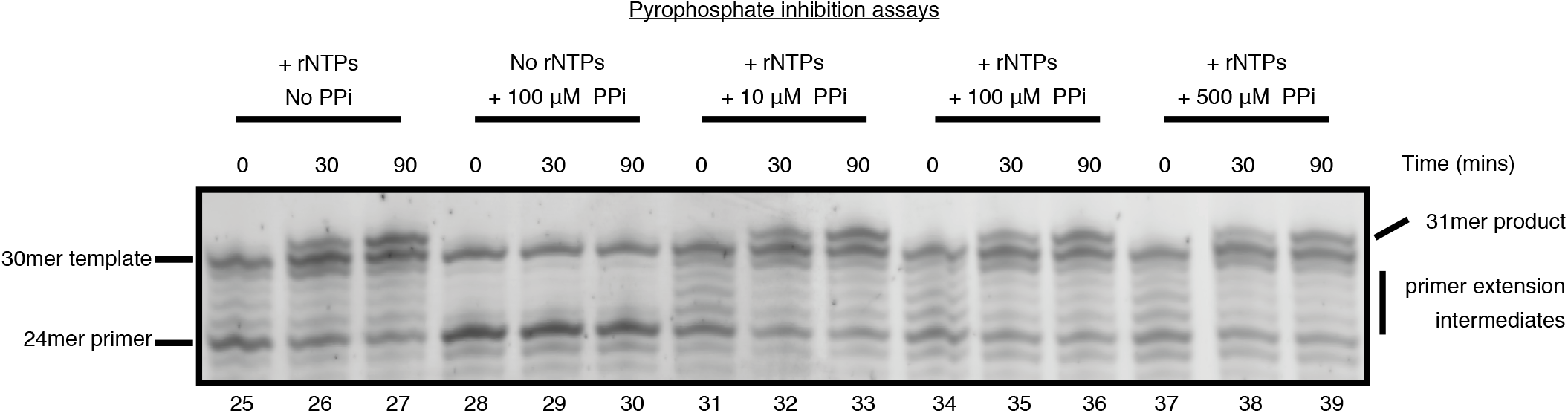
Pyrophosphate inhibition assays of the RNA-dependent RNA polymerase. Primer extension assays by the SARS-CoV-2 RdRp were performed in the presence or absence of rNTPs and/or inorganic pyrophosphate (PPi), and RNA products were resolved by denaturing acrylamide gel electrophoresis and visualized by SYBR Gold staining. The presence of pyrophosphate at concentrations between 10 and 500 μM does not affect the RNA replication by the RdRp.

**Table S1:**
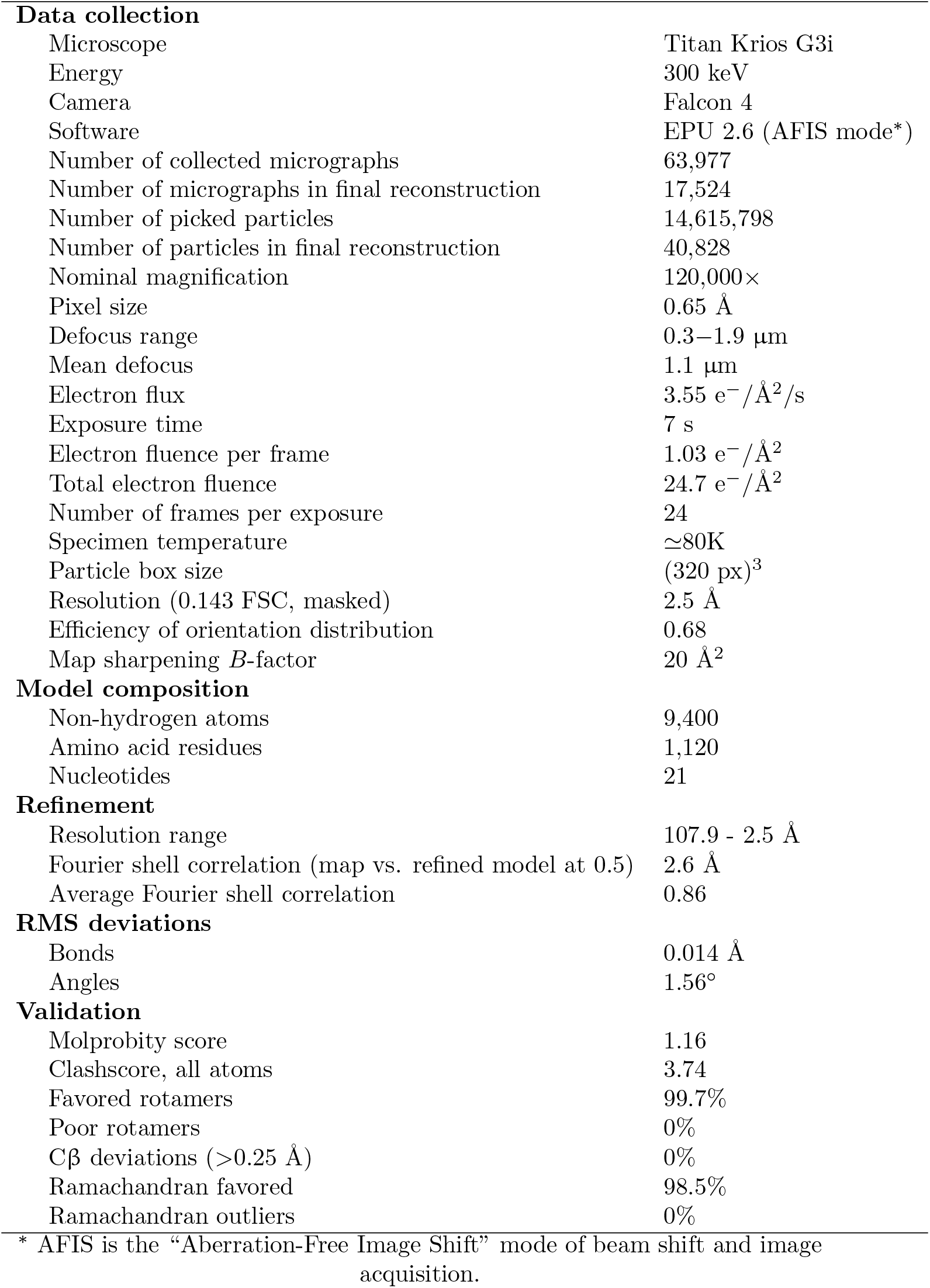
Data, refinement and model statistics.

|  |  |
| --- | --- |
| **Data collection** |  |
| Microscope | Titan Krios G3i |
| Energy | 300 keV |
| Camera | Falcon 4 |
| Software | EPU 2.6 (AFIS mode*) |
| Number of collected micrographs | 63,977 |
| Number of micrographs in final reconstruction | 17,524 |
| Number of picked particles | 14,615,798 |
| Number of particles in final reconstruction | 40,828 |
| Nominal magnification | 120,000× |
| Pixel size | 0.65 Å |
| Defocus range | 0.3–1.9 µm |
| Mean defocus | 1.1 µm |
| Electron flux | 3.55 e<sup>−</sup>/Å<sup>2</sup>/s |
| Exposure time | 7 s |
| Electron fluence per frame | 1.03 e<sup>−</sup>/Å<sup>2</sup> |
| Total electron fluence | 24.7 e<sup>−</sup>/Å<sup>2</sup> |
| Number of frames per exposure | 24 |
| Specimen temperature | ≈80K |
| Particle box size | (320 px)<sup>3</sup> |
| Resolution (0.143 FSC, masked) | 2.5 Å |
| Efficiency of orientation distribution | 0.68 |
| Map sharpening *B*-factor | 20 Å<sup>2</sup> |
| **Model composition** |  |
| Non-hydrogen atoms | 9,400 |
| Amino acid residues | 1,120 |
| Nucleotides | 21 |
| **Refinement** |  |
| Resolution range | 107.9 - 2.5 Å |
| Fourier shell correlation (map vs. refined model at 0.5) | 2.6 Å |
| Average Fourier shell correlation | 0.86 |
| **RMS deviations** |  |
| Bonds | 0.014 Å |
| Angles | 1.56° |
| **Validation** |  |
| Molprobity score | 1.16 |
| Clashscore, all atoms | 3.74 |
| Favored rotamers | 99.7% |
| Poor rotamers | 0% |
| Cβ deviations (>0.25 Å) | 0% |
| Ramachandran favored | 98.5% |
| Ramachandran outliers | 0% |

\* AFIS is the “Aberration-Free Image Shift” mode of beam shift and image acquisition.

